# Individual variation in glucocorticoid plasticity to temperature is not associated with reproductive success in a wild songbird population

**DOI:** 10.64898/2026.09.22.750903

**Authors:** Eve Udino, Caroline Deimel, Michaela Hau, Maria Moiron

## Abstract

Vertebrates cope with rapid temperature fluctuations by plastically adjusting their behaviour and physiology. This phenotypic plasticity is partly mediated by plastic changes in glucocorticoid levels; however, whether individual differences in endocrine plasticity are associated with fitness remains unknown. Here, we examined the relationship between individual variation in glucocorticoid plasticity in response to air temperature changes and reproductive success in a wild great tit population (*Parus major*). Over nine years, we collected repeated measurements of baseline and stress-induced corticosterone during nestling rearing. We found consistent reaction norm patterns in hormone response to temperature variation – as shown in a previous study conducted over a shorter period. This provides robust evidence of individual differences in average hormone levels (intercepts) and the strength of the plastic response (slopes), indicating potential for evolution. However, neither intercepts nor slopes were associated with reproductive success (fledgling number or brood mass), suggesting that individual variation in endocrine plasticity was not under natural selection, at least within the environmental conditions experienced. This study provides a first empirical test of the evolutionary implications of endocrine plasticity in the wild and highlights the need to assess its fitness relevance under more extreme conditions, such as those expected under future climate scenarios.

## Introduction

In the face of global change, temperature alterations pose a major threat to biodiversity (Thomas et al., 2004; Urban, 2015). Beyond increasing mean temperatures, organisms are increasingly confronted with greater thermal variability and a higher frequency and severity of extreme climatic events (Meehl and Tebaldi, 2004; Vasseur et al., 2014; Stillman, 2019; IPCC, 2023). Such thermal alterations have pervasive effects across levels of biological organization, from biochemical and physiological processes to populations and ecosystems. Perhaps the most obvious impacts of gradual temperature changes are shifts in phenology and distribution ranges, whereas extreme thermal events can cause mass-mortality events and breeding failures, as well as impose sublethal costs on individuals (Parmesan, 2006; McKechnie and Wolf, 2010; Scheffers et al., 2016; Marrot et al., 2017; Cunningham et al., 2021; Taff and Shipley, 2023). Consequently, the ability of animals to cope with rapid and unpredictable thermal fluctuations is a key determinant of their persistence in the Anthropocene.

Reversible phenotypic plasticity (or flexibility) – the ability of a single genotype to express varying phenotypes across environmental gradients (Piersma and Drent, 2003; Whitman and Agrawal, 2009) – is often regarded as a rapid mechanism allowing organisms to adapt to spatial and temporal environmental changes (Pigliucci, 2001; Ghalambor et al., 2007; Chevin et al., 2010). Phenotypic plasticity can help individuals coping with thermal changes by adjusting behavioural and physiological traits associated with thermoregulation, including behavioural responses such as microhabitat selection and panting (Xie et al., 2017; Oswald et al., 2019; Udino and Mariette, 2022) as well as acclimation processes (McKechnie, 2008; Boyles et al., 2011; Tattersall et al., 2012). However, the adaptive value of plasticity ultimately depends on its fitness consequences; specifically, plasticity is considered adaptive when phenotypic adjustments translate into increased fitness. Under such circumstances, plasticity may facilitate adaptive evolution provided that both standing phenotypic and genetic variation exist in the underlying traits and that natural selection acts upon this variation (Price et al., 2003; Ghalambor et al., 2007; Gibert et al., 2019). Given the potential influence of plasticity on evolutionary processes, understanding the mechanisms that regulate phenotypic plasticity is therefore essential for predicting population resilience to climate change.

Endocrine systems play a crucial role in coping with environmental changes, including temperature fluctuations (Angelier and Wingfield, 2013; de Bruijn and Romero, 2018; Ruuskanen et al., 2020; Mentesana and Hau, 2022), as they integrate external and internal information to mediate phenotypic responses. In vertebrates, the activation of the hypothalamic-pituitary-adrenal (HPA) axis results in the release of glucocorticoid hormones into the bloodstream (Sapolsky, 2000; Romero and Wingfield, 2015). At baseline levels, these metabolic hormones bind to the high-affinity mineralocorticoid receptors to support physiological processes to meet daily and seasonal energetic demands (Landys et al., 2006; Romero and Wingfield, 2015). At stress-induced levels, these hormones play a different role as part of the so-called stress-response by also binding to the low-affinity glucocorticoid receptors. Indeed, upon exposure to an acute and unpredictable challenge, glucocorticoid levels increase within minutes to reallocate energy toward immediate survival functions (Landys et al., 2006; Romero and Wingfield, 2015). Due to their pleiotropic effects, glucocorticoids are important mediators of phenotypic plasticity (Hau and Goymann, 2015; Hau et al., 2016) and are thus often assumed to be associated with fitness. While evidence supporting that link with fitness is mixed (Breuner et al., 2008; Bonier et al., 2009), most studies have been conducted at the group level, when substantial variation in glucocorticoid levels exists within and between individuals. Because natural selection acts at the individual level, it is necessary to focus on such individual variation to understand the evolutionary implications of endocrine plasticity (Nussey et al., 2007; Williams, 2008; Hau et al., 2016; Taff and Vitousek, 2016).

In recent years, the importance of quantifying endocrine plasticity at the individual level has been increasingly recognized in ecology and evolution. Although most publications are conceptual, they establish important foundations and guidelines in this emergent field (e.g., (Guindre-Parker, 2020; Grindstaff et al., 2022; Malkoc et al., 2022; Taff et al., 2024; Malkoc, 2026). To measure individual variation in endocrine plasticity, a reaction norm approach can be applied. Reaction norms are a statistical tool that allows describing the response of a plastic trait along an environmental gradient such as temperature (Nussey et al., 2007; Martin et al., 2011; Dingemanse and Dochtermann, 2013). Reaction norms decompose the individual response into an intercept – representing the elevation or the average trait at the average environmental variable – and a slope, representing the plasticity of the trait (Malkoc et al., 2022). Thus, applying this approach requires overcoming the challenges of obtaining repeated measurements for each individual along the environmental gradient and gathering a sample size large enough for robust statistical analysis (Malkoc et al., 2022). While reaction norms are commonly used in the field of animal behaviour, such data requirements may explain the lack of empirical studies thus far in endocrinology because individuals need to be repeatedly sampled across the environmental gradient while standardizing other conditions (e.g. reproductive stage). To our knowledge, only three empirical studies have assessed glucocorticoid reaction norms to temperature in birds, experimentally (Baldan et al., 2021) or in the wild (Hau et al., 2022; Taff et al., 2025). In our previous study, wild great tits (*Parus major*) differed in their glucocorticoid plasticity in response to natural temperature variation during the breeding season (Hau et al., 2022). Glucocorticoid levels increased at colder temperatures both at the population and individual levels, however individuals differed in this response to temperature with some birds showing higher elevations or greater plasticity than others (Hau et al., 2022). This increase at colder temperatures was also found in the study of a large database including different birds species sampled across the world, with different response patterns between species (Taff et al., 2025). These studies provide evidence of standing phenotypic variation in endocrine plasticity, a prerequisite for evolutionary change. However, whether this variation is a target of natural selection and therefore a potential evolutionary response to climate change remains to be determined.

Thus, in this study, we examined the relationship between individual differences in corticosterone plasticity (the main avian glucocorticoid) in response to environmental temperature and reproductive success in wild great tits. Following from our previous five-year study that established individual differences in corticosterone reaction norms to temperature (Hau et al., 2022), we continued studying this population to determine the fitness consequences of this endocrine plasticity. Briefly, from 2015 to 2023, repeated measurements of baseline and stress-induced corticosterone as well as reproductive success metrics were obtained across years in adults during the nestling rearing stage. First, we tested the robustness of our previous findings in corticosterone plasticity to temperature in this 9-year dataset by running a series of univariate mixed models. Second, we assessed the adaptive value of this corticosterone plasticity (adaptive, maladaptive, neutral), by using bivariate mixed models to test the covariance between the reaction norm components (elevation and plasticity) of baseline or stress-induced corticosterone and two reproductive success metrics, measured as fledgling number or brood mass. For both baseline and stress-induced corticosterone reaction norms to temperature, we hypothesized that if a greater hormonal plasticity (i.e. steeper slope) reflects a greater ability to adjust to environmental changes, it may confer fitness benefits. We therefore predicted that (i) individuals with steeper slopes would have a higher reproductive success, that is, a higher fledgling number and/or brood mass (i.e., positive covariation). We also hypothesized that, if environmental conditions increase physiological and energetic demands, then individuals have to increase resources allocation towards self-maintenance at the expense of reproduction. We therefore predicted that (ii) individuals with a higher intercept (i.e., more resources allocated toward self-maintenance) would have a lower fledgling number and/or brood mass (i.e., negative covariation).

## Methods

### Study population, breeding monitoring and reproductive success

The study was carried out in a free-living nest box population of great tits located in the Ettenhofer Holz, (Upper Bavaria, Germany, 48°03’27.8”N 11°15’15.6”E), a mixed managed forest, during the breeding seasons from 2015 to 2023. From April to July, active nest boxes were monitored daily (or every two or three days in case of inclement weather) during the egg laying period until the first day of incubation, from day 11 of incubation until hatching (day 0), and from day 15 of nestling rearing until fledgling to record dates of phenological events and number of eggs and nestlings.

A nest was considered successful if at least one chick fledged. The fledgling number was calculated as number of nestlings at day 15 minus dead nestlings found in the nest after fledgling (in all nests, median: 4.0, range: 0 – 9; in successful nests only, median: 5.0, range: 1 – 9). On day 15, nestlings were weighed to calculate the brood mass, i.e. the sum of all nestling masses (in successful nests only, median: 80.75g, range: 16.30 – 158.75g).

### Capture and blood sampling

Breeding pairs were caught when the nestlings were 8 days old, or between 7 and 11 days old in case of inclement weather. If one of the parents could not be caught on capture day, its capture was attempted within the next two days. Birds were caught individually upon entering the nest box, using a remote-controlled or mechanical trap, or by blocking the entrance hole with a bird bag (Ecotone). All captures occurred between 7:00 and 14:00 to minimise the effect of diel corticosterone variation.

A first blood sample was collected within 3 minutes of capture by puncturing the ulnar vein with a 26-gauge needle and drawing blood with a heparinised microcapillary tube (baseline corticosterone sample, C0; bleeding time mean: 2.05 minutes, range: 55 seconds – 4.00 minutes with 97.4% of samples < 3 minutes). Once the bleeding was stopped, the bird was kept in an opaque cotton bird bag as part of a capture-restraint protocol. The bird was then handled to record its identity (or band it) and its morphometric measurements (including body mass) before the collection of a second blood sample 30 minutes after capture (stress-induced corticosterone sample, C30; bleeding time: mean: 30.08 minutes, range: 19.75 – 45 minutes). Once the bleeding stopped, the bird was returned to the nest box via the entrance hole. If the condition of a bird after catching was not satisfying or deteriorated after the first blood sample, no sample was collected and the bird was released after taking its morphometric measurements. This included low body mass, reduced breast muscle, behavioural signs of distress (e.g., body tone, unresponsiveness, closed eyes) or increased blood loss.

Blood samples were kept on ice for up to 7 hours until centrifugation in the laboratory at 2000g and 4°C for 10 minutes to separate plasma and blood cells. Plasma samples were transferred into separate tubes and stored at −20°C until corticosterone levels were assayed.

### Air temperature at capture

Among different temperature variables (mean, minimum, maximum of capture day, mean of previous or three previous days), both baseline and stress-induced corticosterone levels were best explained by the air temperature at capture (Hau et al., 2022), hence only this variable was used in this study (median: 13.6°C, range: 4 – 27°C).

The temperature at capture was noted at the time of the baseline blood sample, using a thermometer (BASETech Thermometer E0217, accuracy ±1°C). When the temperature could not be recorded, the temperature from the weather station was used instead (HOBO Micro Station Data Logger H21-002 fitted with S-THB-M002 temperature and relative humidity sensor, accuracy ±0.21°C). The weather station is located at the centre of the study area and recorded data every 30 minutes. Therefore, the temperature record closest to the time of capture was used. Temperatures from the thermometer and the weather station were previously found to be highly correlated (Hau et al., 2022).

### Corticosterone assays

Corticosterone was extracted from plasma samples following a double liquid-liquid extraction procedure with diethyl ether. Samples were dried under nitrogen gas and reconstituted with up to 240µL assay buffer, with dilutions ranging from 1:10-1:30 and 1:30-1:96 for baseline and stress-induced samples respectively. The reconstituted samples were stored overnight at 4°C and assayed the next day in duplicate using commercial Enzyme-Linked Immunosorbent Assays (ELISA) kits, from Enzo Life Sciences (ADI-900-097) in 2015, and Arbor Assays (K014-H5) from 2016 to 2023. Further details on the assays, controls and coefficients of variation are available in the supplementary materials; details on assay validations and hormone extraction are available in the supplementary materials of (Hau et al., 2022).

### Sample sizes

In total, 829 captures were made, with 61 to 113 captures per year. This corresponds to a total of 435 individuals caught, including 210 males and 225 females. Among these birds, one or repeated measurement(s) of corticosterone levels were collected, mostly across successive years except for birds with second clutches. For baseline corticosterone, 209 individuals had one measurement, 113 had two, 53 had three, 26 had four, 12 had five, 3 had six and 1 had seven measurements, for a total of 783 samples. For stress-induced corticosterone, 217 individuals had one measurement, 116 had two, 47 had three, 29 had four, 12 had five, 1 had six and 1 had seven measurements, for a total of 779 samples. Individuals with single corticosterone measurements were included in the analyses because these single data increase statistical power while reducing credible intervals around the variance (Martin et al., 2011).

### Statistical analyses

All statistical analyses were performed with R (R Core Team, 2024, v4.4.1) and RStudio (v2024.04.2).

### Individual variation in corticosterone plasticity in response to temperature

First, we aimed to test the robustness of glucocorticoid reaction norm patterns described in our previous study (Hau et al., 2022) on this extended version of the dataset. Therefore, we repeated the analyses consisting in four models, with the difference that the variable *year* was used as continuous variable in our analysis.

Briefly, model 1 assesses the population response to temperature, fitting corticosterone levels (baseline, C0; or stress-induced, C30) as function of *sex, year, bleeding time*, and *air temperature at capture*, with bird identity included as random factor. In all models, corticosterone levels (C0 and C30) were fitted with a Gaussian error distribution.

Model 2 tests the contributions of between- and within-individual responses to temperature, following a “within-subject centring” approach (van de Pol and Wright, 2009). To do so, we calculated the *between-individual component* as the average temperature experienced by each individual across its repeated captures, and the *within-individual component as* the temperature deviation for each capture from this individual average. Then, the model fitted corticosterone levels (C0 or C30) as function of the *sex, year, bleeding time* and the *between*- and *within-individual effects of temperature*, with *bird identity* included as random factor. Lastly, we tested whether the between- and within-individual effects of temperature differed by calculating the difference in their posterior means and estimating their 95% credible intervals.

Model 3 assesses between-individual differences in their plastic corticosterone response to temperature. In this random regression mixed model, corticosterone levels (C0 or C30) were fitted as response variable, and the same fixed effect structure as model 1 was used. *Bird identity* was included as random intercept and *air temperature at capture* was included as random slope. This model was run twice, assuming residual homogeneity or heterogeneity across years (i.e., the model allowed residual variation to differ among the nine years of sampling). Two other models were run including random intercepts only and homogeneous or heterogeneous residuals, and the four models were compared using deviance information criterions (DIC; lower values indicating better model fit).

Model 4 tested the covariation between baseline (C0) and stress-induced (C30) corticosterone levels. A bivariate model was fitted, including *C0* and *C30* corticosterone levels as response variables, the same fixed and random structures as model 1, and using an unstructured covariance matrix.

### Fitness consequences of corticosterone plasticity

Second, we aimed to test whether glucocorticoid reaction norms and reproductive success covary. A bivariate model was fitted (model 5), including either *baseline* (C0) or *stress-induced* (C30) corticosterone levels and *fledgling number* as response variables. Corticosterone levels were fitted with a Gaussian distribution and fledgling number with a zero-inflated Poisson error distribution with a fixed residual variance for the zero-inflated component of five. This zero-inflated distribution includes a Poisson component that models fledgling number as a count, and a zero-inflated component that models fledgling number as the probability of excess zeros, corresponding here to nest failures (i.e., breeding attempts resulting in zero fledglings). *Sex* and *year* were included as fixed effects for both variables, and *air temperature at capture* and *bleeding time* were included as fixed effects only for corticosterone levels. Since the random regression mixed model was the best fit to the data (model comparisons from model 3, *bird identity* was included as random intercept and *air temperature* as random slope only for corticosterone levels. This model was run three times, (model 5a) in all data and only in (model 5b) males or (model 5c) females to uncouple the breeding mates, sharing the same reproductive success values but possibly different hormone profiles, which could hinder a covariation between the two response variables.

Given the complexity associated with the use of the zero-inflated Poisson distribution in model 5 above in terms of computing and interpretation of results, we also present an alternative analysis using two models. The first model, model 6, focusses on the binomial nest outcome (success or failure), fitting *corticosterone levels* (C0 or C30) and *nest outcome* as response variables, and *sex, year, bleeding time*, and *air temperature at capture* as fixed effects. Corticosterone levels were fitted using a Gaussian error distribution, and reproductive outcome using a binomial error distribution with a logit link function and fixed the residual variance to one. The second model, model 7, is similar, but uses *fledgling number* as response variable for reproductive success and was run with data from successful nests only to obtain satisfying convergence of the model. Fledgling number was fitted using a Poisson error distribution. Lastly, the body condition, or quality, of great tit fledglings may also be important for their survival (Naef-Daenzer et al., 2001; Naef-Daenzer and Grüebler, 2016 but see Jones et al., 2026), which would increase the fitness of the parents. Therefore, we ran model 8, similar to model 7 but replaced fledgling number by *brood mass*, which integrates both fledgling quantity and quality. Brood mass was fitted with a Gaussian error distribution.

All continuous fixed effects were scaled. All Bayesian models were fitted using the package MCMCglmm (Hadfield, 2010) and parameter-expanded priors. The number of iterations, thinning interval and burn-in were set to obtain an effective sample size of 1000 and autocorrelation values below 0.1. The convergence of the Markov chain Monte Carlo was assessed using Heidelberger-Welch diagnostic and by visual inspection of the posterior draws. To describe posterior distributions, we report posterior modes, medians (less sensitive to skewed distributions than the mean) and 95% credible intervals (Pick et al., 2023). We consider effects to be relevant, or meaningful, when their 95% credible intervals do not overlap zero.

## Results

### Individual variation in corticosterone plasticity along temperature

Overall, we observed patterns of corticosterone plasticity in relation to air temperature at capture that are consistent with those previously reported in the same population over a shorter study period (Hau et al., 2022), confirming the robustness of these findings. Briefly, the different univariate models show evidence of hormonal plasticity in response to temperature, with both baseline and stress-induced corticosterone levels increasing at lower temperature at the population [posterior median (95% CI) for baseline levels: −0.446 (−0.627, −0.257), stress-induced levels: −2.303 (−3.132, − 1.572); model 1; table S1] and at the individual level [“Within-ID temperature effect” for baseline levels: −0.108 (−0.175, −0.054), stress-induced levels: −0.521, (−0.782, −0.276); model 2; table S2]. In addition, there is evidence of between-individual variation in elevation (i.e., intercept) and plasticity (i.e., slope) in both hormonal traits, indicating that some individuals had higher average corticosterone levels than others and that some individuals had a greater plasticity in response to temperature than others [“Individual intercept” for baseline levels: 1.035 (0.492, 1.698), stress-induced levels: 36.964 (23.81, 48.978); “Individual slope” for baseline levels: 0.281 (0, 0.62), stress-induced levels: 0.479 (0, 4.09); model 3; table S4]. All model results are fully described in the supplementary material.

### Sources of phenotypic variation in corticosterone and reproductive success

The bivariate analysis shows that both baseline and stress-induced corticosterone levels increased at lower air temperature at capture (model 5a, table 1). In addition, baseline levels decreased through the years whereas stress-induced levels increased (model 5a, table 1, figure S1). Baseline levels increased with longer latency to collect the blood sample (“bleeding time”) as expected, and were also higher in females, while these effects were absent in stress-induced levels (model 5a, table 1). To some extent, for both hormonal traits, the Poisson component of fledgling number increased weakly across years (model 5, table 1). However, the zero-inflated component of fledgling number showed no variation across years, indicating no temporal change in the occurrence of nest failures over time (model 5a, table 1).

**Table 1.** Model 5a: summary statistics of a bivariate mixed model to test for the covariance and correlation between baseline or stress-induced corticosterone levels and fledgling number in the full dataset. Hormonal levels were fitted using a Gaussian error distribution and reproductive success using a zero-inflated Poisson distribution. Therefore, reproductive success shows estimates for the Poisson and zero-inflated (ZI) part of the distribution. Effect sizes are considered relevant when their 95% credible intervals (CI) do not overlap zero. Variables of interest for this model are in bold. Baseline corticosterone: N = 760 samples from 406 birds; Stress-induced corticosterone: N = 743 samples from 408 birds.

|  |  | Baseline corticosterone |  |  | Stress-induced corticosterone |  |  |
| --- | --- | --- | --- | --- | --- | --- | --- |
|  |  | Posterior mode | Posterior median | 95% CI | Posterior mode | Posterior median | 95% CI |
| Fixed effects | Response |  |  |  |  |  |  |
| Intercept | CORT | 4.371 | 4.374 | [4.067, 4.677] | 25.498 | 25.477 | [24.205, 26.712] |
|  | Fledglings | 1.548 | 1.555 | [1.503, 1.62] | 1.56 | 1.559 | [1.502, 1.617] |
|  | Fledglings (ZI) | -2.953 | -3.004 | [-3.696, -2.235] | -2.89 | -3.043 | [-3.836, -2.423] |
| Sex (female) | CORT | 0.716 | 0.624 | [0.225, 1.067] | 1.065 | 1.069 | [-0.771, 2.823] |
|  | Fledglings | -0.023 | -0.029 | [-0.111, 0.047] | -0.01 | -0.027 | [-0.109, 0.057] |
|  | Fledglings (ZI) | 0.281 | 0.306 | [-0.632, 0.985] | 0.021 | 0.114 | [-0.6, 0.948] |
| Year | CORT | -0.496 | -0.51 | [-0.731, -0.324] | 1.148 | 1.408 | [0.554, 2.298] |
|  | Fledglings | 0.034 | 0.036 | [-0.001, 0.079] | 0.044 | 0.036 | [-0.005, 0.072] |
|  | Fledglings (ZI) | -0.208 | -0.202 | [-0.581, 0.153] | -0.184 | -0.236 | [-0.618, 0.144] |
| Bleeding time | CORT | 1.033 | 1.02 | [0.821, 1.19] | -0.157 | -0.023 | [-0.85, 0.774] |
| Air temperature | CORT | -0.449 | -0.481 | [-0.686, -0.274] | -2.744 | -2.548 | [-3.401, -1.779] |
| **Random effects** |  |  |  |  |  |  |  |
| Intercept variance | CORT | 1.172 | 1.137 | [0.456, 1.852] | 32.824 | 31.351 | [17.937, 47.963] |
|  | Fledglings | 0 | 0.001 | [0, 0.006] | 0 | 0.001 | [0, 0.005] |
|  | Fledglings (ZI) | 3.176 | 2.871 | [0, 6.305] | 2.468 | 3.033 | [0.003, 6.774] |
| Slope variance | CORT slope | 0.3 | 0.413 | [0.035, 0.91] | 0.029 | 0.87 | [0, 4.806] |
| Intercept-intercept cov. | **CORT-fledglings** | 0 | 0.014 | [-0.018, 0.076] | -0.001 | 0.013 | [-0.166, 0.236] |
| Intercept-intercept corr. |  | 0.905 | 0.617 | [-0.764, 0.999] | 0.585 | 0.198 | [-0.803, 0.957] |
| Intercept-intercept cov. | **CORT-fledglings (ZI)** | 0.227 | 0.353 | [-0.396, 1.116] | -3.094 | -3.154 | [-7.678, 0.087] |
| Intercept-intercept corr. |  | 0.201 | 0.206 | [-0.3, 0.799] | -0.328 | -0.35 | [-0.781, 0.052] |
| Intercept-intercept cov. | Fledglings-fledglings (ZI) | 0 | 0.001 | [-0.057, 0.064] | 0 | -0.005 | [-0.113, 0.048] |
| Intercept-intercept corr. |  | 0.145 | 0.057 | [-0.815, 0.831] | -0.854 | -0.26 | [-0.98, 0.862] |
| Intercept-slope cov. | CORT slope-CORT | -0.507 | -0.566 | [-1.014, -0.176] | -0.252 | -1.766 | [-7.155, 2.256] |
| Intercept-slope corr. |  | -0.954 | -0.87 | [-0.999, -0.548] | -0.506 | -0.386 | [-0.984, 0.529] |
| Intercept-slope cov. | **CORT slope-fledglings** | 0 | -0.007 | [-0.046, 0.017] | 0 | -0.001 | [-0.066, 0.037] |
| Intercept-slope corr. |  | -0.84 | -0.57 | [-0.999, 0.808] | -0.758 | -0.174 | [-0.95, 0.912] |
| Intercept-slope cov. | **CORT slope-fledglings (ZI)** | -0.193 | -0.174 | [-0.804, 0.37] | -0.004 | 0.673 | [-0.785, 3.161] |
| Intercept-slope corr. |  | -0.231 | -0.185 | [-0.89, 0.362] | 0.804 | 0.519 | [-0.523, 0.998] |
| Residual (within-ID) variance | CORT | 5.766 | 5.954 | [5.151, 6.903] | 96.178 | 97.814 | [83.435, 111.363] |
|  | Fledglings | 0.03 | 0.03 | [0.021, 0.041] | 0.031 | 0.03 | [0.021, 0.042] |
|  | Fledglings (ZI) | 5 | 5 | [5, 5] | 5 | 5 | [5, 5] |

### Fitness consequences of corticosterone plasticity

Contrary to our predictions, for both baseline and stress-induced corticosterone levels, the bivariate analysis shows no evidence of covariation between either the elevation or plasticity of hormonal levels and both the Poisson and zero-inflated components of fledgling number (“Intercept-intercept cov.”, “Intercept-slope cov.”, model 5a, table 1 figure 1). This translated into a subsequent lack of support of correlation between those parameters, as we note the large uncertainty in the credible intervals (“Intercept-intercept corr.”, “Intercept-slope corr.”, model 5a, table 1, figure 1). The results were consistent when running this model separately for males and females (models 5b-5c, tables S6-S7).

**Figure 1.**
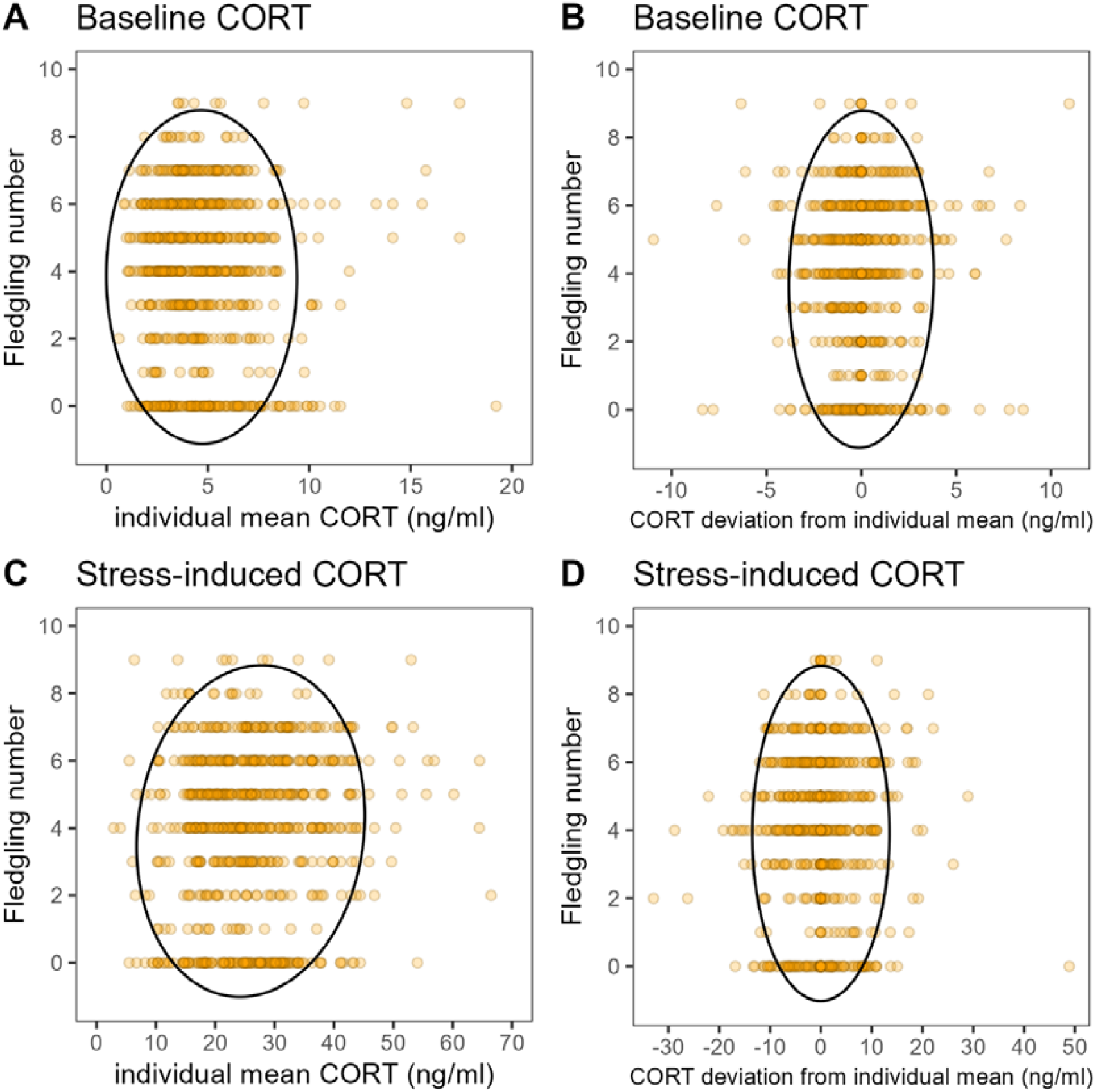
Fledgling number along individual corticosterone parameters at baseline (A, B) and stress-induced (C, D) levels. Corticosterone parameters consist in the individual mean across repeated measurements (A, C) and deviation from this mean for each measurement (B, D). The solid black ellipse represents the variance in fledging success, variance in corticosterone parameters, and their covariance, providing visual representation of the joint variability of the two variables. Note that this figure is drawn for illustrative purposes only, conclusions were drawn directly from the model estimates (Table 1).

This finding is corroborated by the two supplementary models that we performed. Neither the elevation nor the plasticity of any hormonal trait covaried with nest outcome as success or failure [“Int.-int. cov.”: posterior median (95% CI) for baseline levels: 0.214 (−0.296, 0.671), stress-induced levels: −2.028 (−4.662, 0.181); “Int.-slope cov.”: baseline levels: −0.099 (−0.455, 0.258), stress-induced levels: 0.338 (−0.364, 1.973), model 6, table S8] or, after restricting the analysis to successful nests only, with fledgling number (“Int.-int. cov.”: baseline levels: 0.01 (−0.021, 0.068), stress-induced levels: 0.011 (−0.125, 0.261), “Int.-slope cov.”: baseline levels: −0.005 (−0.041, 0.014), stress-induced levels: 0 (−0.047, 0.037), model 7, table S9) or brood mass (“Int.-int. cov.”: baseline levels: 3.491 (−0.555, 9.289), stress-induced levels: 7.436 (−14.306, 31.065), “Int.-slope cov.”: baseline levels: −1.45 (−5.508, 0.988), stress-induced levels: −1.069 (−11.796, 8.895), model 8, table 10) as reproductive success metrics.

## Discussion

In this 9-year study in wild great tits we investigated whether individual variation in glucocorticoid plasticity in response to varying air temperature was under selection. Although we observed clear evidence for individual differences in average expression (intercepts) and plastic responses (slopes), this variation in reaction norms was, against our prediction, not associated with variation in two different reproductive success metrics. Thus, we found no detectable evidence that glucocorticoid plasticity was associated with fitness benefits or costs, suggesting that, in our population, this plasticity may be neutral, or have limited adaptive or maladaptive consequences. To our knowledge, this is the first empirical study to test the adaptive significance of glucocorticoid plasticity in a wild avian population. The absence of selection on endocrine plasticity in this study raises fundamental questions about the evolutionary implications of hormonal plasticity in the context of climate change and highlights the need for further work.

As expected, baseline and stress-induced corticosterone levels increased at lower air temperature at capture at the population level. Corticosterone levels also increased at lower temperature within individuals, and we found important between-individual differences in average values and plastic responses. These results are consistent with those previously reported in our shorter-term study period (Hau et al., 2022); providing robustness towards the findings. The main objective of the current study was to extend the scope of the previous analyses and investigate whether such differences in corticosterone reaction norms were selected for, and hence, associated with reproductive success.

Surprisingly, neither the elevation nor the slope of the corticosterone reaction norms was associated with fledgling number or brood mass for both baseline and stress-induced levels. This result was maintained even when analyses were separated by sex. This main finding suggests that, in this population, natural selection did not act in a detectable manner on corticosterone plasticity to temperature variation, a key natural gradient for this species. Although no previous study has ever tested the relationship between an individual’s endocrine plasticity and its reproductive success, theory expects glucocorticoid plasticity to be adaptive, and possibly under different selective pressures given the distinct roles of baseline and stress-induced glucocorticoids (Vitousek et al., 2019 but see (Béziers et al., 2019). Nonetheless, the association between circulating glucocorticoid levels and reproductive success has been the focus of many studies, and evidence supporting such relationship remains mixed. A meta-analysis shows that both baseline and stress-induced levels correlated negatively with reproductive success across vertebrate taxa (Schoenle et al., 2021), but this pattern is often absent or reversed in individual studies (reviewed in (Bonier et al., 2009; Hau et al., 2016), but see also (Patterson et al., 2014; Vitousek et al., 2018). Such complexity is often explained by the context dependence of hormonal responses (Schoenle et al., 2018), such as life-history stage or seasonality, variables used to measure reproductive success, or other physiological traits accounted for (like antioxidants (Mentesana et al., 2024). We note however that in great tits, individuals with high baseline corticosterone before breeding and low levels during breeding had the highest reproductive success. (Ouyang et al., 2013). The closest study to investigate the adaptive significance of endocrine plasticity was conducted in primates, were faecal glucocorticoids were associated with survival (Carrera et al., 2025).

Multiple, non-mutually exclusive explanations could account for the lack of an association between endocrine plasticity and reproductive success in our study. First, from a biological perspective, selection on corticosterone plasticity may have been absent because the environmental conditions experienced by our study population were generally not sufficiently challenging. Our study population, located in a temperate climate of western Europe, experienced relatively mild temperatures at capture ranging between 4-27°C. The thermoneutral zone of great tit ranges between 15-30°C in a Swedish population (Broggi et al., 2005), and 17-35°C in a Spanish population (Playà-Montmany et al., 2021). Thus, captures occurred both within and below this range; specifically, 61% of captures occurred below 15°C, but only 15% occurred below 10°C, where thermal conditions imposed a greater challenge. This lower proportion can be explained both by a lower occurrence of these lower temperatures at the time of captures, but also because, for ethical concerns, catching at these temperatures were generally avoided when possible to prevent triggering nest abandonment. Nevertheless, metabolic rate shows a continuous and marked increase below 15°C (Malkoc et al., 2021), indicating that birds captured at 10–15°C still have experienced increased energetic demands. Overall, most individuals experienced mild thermal conditions within or close to the thermoneutral zone, where metabolic demands are lowest and temperature-dependent corticosterone responses limited. For instance, in tree swallows (*Tachycineta bicolor*), baseline corticosterone levels were negatively associated with fledgling number only during a challenging year, but not during milder years (Vitousek et al., 2018). Therefore, it is plausible that the mild environmental conditions experienced by individuals in our study did not impose a sufficiently strong selective pressure for differences in plasticity to translate into fitness differences. Alternatively, the breadth of variation in corticosterone plasticity expressed in our population may have been insufficient for selection to act upon it (Taff et al., 2024). Therefore, our results suggest that the fitness relevance of glucocorticoid plasticity may be conditional on environmental severity, which encourages for further testing in more extreme or unpredictable environments.

Second, the absence of a relationship between corticosterone plasticity and reproductive success could be explained by methodological reasons. For instance, despite nine years of data, our power to detect moderate or weak relationships may not be sufficient. Furthermore, our analysis assumed that corticosterone reaction norms to temperature were linear and temporally stable within our yearly sampling intervals. Given that corticosterone is a metabolic hormone (Jimeno and Verhulst, 2023), its response to temperature is likely non-linear, with sharp increases only below thermoneutrality—a pattern observed across 99 bird species (Taff et al., 2025). Moreover, due to changes in environmental conditions or internal factors, reaction norms may also themselves be plastic within individuals over time and across contexts (Dupont et al., 2024; Malkoc, 2026). Although the temperature range in our study was within and below the thermoneutral zone of the great tits, that is, avoiding a potential U-shaped response over a gradient extended to higher temperatures, these important aspects were not integrated in our models given the present difficulty to fit a model to the data using linear reaction norms. Future studies could explore non-linear relationships, use finer temporal resolution and integrate other behavioural and physiological covariates, but this would require a very large dataset and/or adapted study protocol.

The absence of selection on corticosterone plasticity within mild conditions raises the critical question of evolutionary implications for the population’s adaptive potential in the face of current climate change. In this great tit population, standing variation in corticosterone plasticity within- and between-individuals exists, indicating a potential for selection to act upon this variation in the event of changing environmental conditions. However, for an evolutionary change to occur, endocrine plasticity must also be both under natural selection and heritable. Although we found no evidence of selection in the mild conditions experienced by our population, a moderate heritability of both baseline and stress-induced corticosterone levels was found in other avian species (Jenkins et al., 2014; Stedman et al., 2017; Béziers et al., 2019). In addition, a recent study found a biologically-meaningful heritability in corticosterone plasticity in house sparrows (*Passer domesticus*) within artificial selection settings (Ouyang and Lendvai, 2026), while such heritability in wild population remains untested. Thus, should environmental conditions become more challenging, including extreme climatic events or unpredictable temperature fluctuations, selection may act upon individual variation in this plasticity, fulfilling the necessary conditions for evolutionary change and turning endocrine plasticity into a key component of adaptive capacity.

## Conclusion and future directions

Our 9-year study in wild great tits reveals the presence of substantial individual variation in glucocorticoid plasticity (or flexibility) to air temperature, but contrary to our expectations, this individual variation was not linked with reproductive success under mild environmental conditions. This unexpected result suggests that natural selection did not act on individual variation in glucocorticoid plasticity in this population. Nevertheless, this finding underscores the critical need to examine the fitness consequences of endocrine plasticity across broader environmental gradients and during extreme conditions, such as those expected under future climate change projections. Future studies could investigate whether selection on glucocorticoid plasticity emerges only during periods of thermal challenges (e.g., heatwaves or cold snaps) or climatic unpredictability, which may reveal context-dependent fitness benefits. Furthermore, future research could move beyond circulating hormone levels to focus on other components involved in endocrine plasticity, such as response speed, negative feedback (Zimmer et al., 2019), receptor or protein expression (Hofmeister and Rubenstein, 2016; Zimmer et al., 2023, 2024; Jimeno and Rubalcaba, 2024). Moreover, that hormonal plasticity did not result in observed benefits on reproductive success also raises questions from a physiological perspective. Notably, future research could assess the costs of maintaining a plastic phenotype in endocrine levels, such as energetic trade-offs or components of oxidative stress. Given the pleiotropic role of glucocorticoids in the organism, understanding how endocrine plasticity integrates with other physiological traits and behaviour would be an interesting avenue for further work (“plasticity syndromes”, Malkoc, 2026). Despite the methodological and statistical challenges inherent to reaction norm and long-term field studies, empirical work is needed not only to advance our understanding of the adaptive capacity of populations in a changing world, but also to inspire other fields and policy makers to acknowledge individual variation in plastic responses as a crucial but underappreciated biological trait that is meaningful beyond evolutionary science.

## Ethics

The procedures involving animals were conducted in accordance with the German legal requirements and were approved by the regional governmental authority, the Regierungspräsidium von Oberbayern (permit no. ROB-55.2-2532-Vet_02-13-204, ROB-55.2-2532-Vet_02-15-25, and ROB-55.2-2532.Vet_02-19-51).

## Data accessibility

The dataset and R code will be made available on an online repository upon acceptation to publication.

## Authors’ contributions

**EU:** Conceptualization, Data Curation, Formal Analysis, Investigation, Methodology, Software, Validation, Visualization, Writing – Original Draft Preparation, Writing – Review & Editing; **CD:** Data Curation, Investigation, Validation, Writing – Review & Editing; **MH:** Conceptualization, Funding Acquisition, Methodology, Project Administration, Supervision, Validation; **MM:** Formal Analysis, Methodology, Software, Supervision, Validation, Visualization, Writing – Original Draft Preparation, Writing – Review & Editing

## Conflict of interest

We declare we have no competing interests.

## Funding

This study was funded by the Max Planck Institute for Biological Intelligence. EU was supported by the Alexander von Humboldt foundation, and MM by the German Research Foundation (grant number 540199894).

## Acknowledgements

We dedicate this contribution to the memory of Michaela Hau. Ela started this long-term field study to investigate the evolutionary drivers that shape endocrine systems, and in this contribution, we finally could investigate “THE” question she was really curious to find an answer to. Unfortunately, Ela passed away before the analyses were finished, but she saw preliminary findings. Her keen understanding of endocrinology and evolution, her deep insights and thoughtful comments and her warm, funny, and determined personality made this study possible and are deeply missed. She was a leader in the field of evolutionary physiology, and she continues to inspire us.

Long-term field studies rely on the contributions of many people to produce a unique dataset, by collecting data in the field and running analyses in the lab. We want to thank all the contributors of this study for their effort, passion, and dedication: Nataly Hidalgo Aranzamendi, Carlotta Bonaldi, Robert de Bruijn, Skylar Buckingham, Julia Cramer, Tanguy Deville, Holland Galante, Sam Hardman, Sabine Jörg, Saverio Lubrano, Natalia Perez-Ruiz and Michael Stiegler. We are grateful to the Erzdiozöse München und Freising, particularly to K. Meindl, B. Vollmar and M. Laußer, for allowing us to work in their forest. We also thank the Hau research group for providing feedback over the course of the study, with a special thanks to Kasja Malkoc for useful comments on the manuscript draft.

## Notes

### Competing Interest Statement

The authors have declared no competing interest.

