## supplementary material for "Individual variation in glucocorticoid plasticity to temperature is not associated with reproductive success in a wild songbird population"

### 1. Methods

**Corticosterone assays**

Within years, all samples from one individual were run on the same plate. The plates were read at 450nm using a microplate spectrophotometer (iMark™ 17172, Bio-Rad), sample corticosterone concentrations were calculated using a four-parameter standard curve fit and corrected by the dilution factor.

Two negative blank controls (ROTISOLV® HPLC grade water, A511.1, Carl Roth) were added as first and last samples of each hormone extraction batch and run on the plate to detect potential contamination. Positive controls were also extracted and run in duplicate to calculate intra- and inter-plate coefficients of variation (CV). They consisted of stripped chicken plasma into which a known quantity of corticosterone standard (100 000 pg/mL, provided with the kits) was added to obtain the desired final concentration. These controls were run as two duplicates, added at the beginning and the end of the samples to capture variation across the plate due to pipetting.

For Enzo assays, one quality control was used, at 52% binding, corresponding to a corticosterone concentration of 634 pg/mL. Across 8 plates, the average intra-plate CV was 13% and the inter-plate CV was 16%. For Arbor Assays plates, two quality controls were used, at 69% (174 pg/mL) and 40% binding (736 pg/mL) to fall in the range of baseline and stress-induced concentrations respectively. Across 59 plates, the average intra-plate CVs are 11% at 69% binding and 5% at 40% binding, and the inter-plate CVs are 17% at 69% binding and 9% at 40% binding. CVs were calculated using the quality control sample concentration. Note that the inter-plate CVs also capture variation in extraction efficiency across plates. Across all 2825 samples assayed, the average CV between duplicates was 2.1% (calculated from the percentage of binding values).

Sample measurements that were below the detection limit of the plate they were measured on were replaced with the respective plate's detection limit, calculated as the mean optical density of the maximum binding (Zero, B0) duplicate wells minus two times the standard deviation between the duplicates.

A detailed description of the methods can be found in the supplementary materials of Hau M, Deimel C, Moiron M. 2022. Great tits differ in glucocorticoid plasticity in response to spring temperature. Figshare. (https://doi.org/10.6084/m9.figshare.c.6261884).

### 2. Results

At the population level, both baseline and stress-induced corticosterone levels increased at lower temperatures (model 1, table S1). Interestingly, baseline corticosterone levels decreased through the years whereas stress-induced levels increased (model 1, table S1, figure S1). In addition, for baseline corticosterone levels only, hormonal levels were higher in females than in males (model 1, table S1), and increased with a longer bleeding time (latency to collect the blood sample; model 1, table S1).

At the individual level, both baseline and stress-induced corticosterone levels showed within-individual variation in response to temperature ("Within-ID temperature effect", model 2, table S2) as well as between-individual variation ("Between-ID temperature effect", model 2, table S2). There was no difference between these within- and between-individual effects ("Difference of between-within ID effects", model 2, table S2).

Both hormonal traits also showed between-individual variation in reaction norm components. Among the four types of model, the model comparison revealed that the best fitting model included a random intercept for bird identity, a random slope for temperature and a heterogeneous residual variance structure across years (table S3). In this model, for both baseline and stress-induced corticosterone levels, results indicate between-individual variation in elevation (ie, intercept) and in plasticity (ie, slope) in response to temperature (model 3, table S4). In addition, for baseline corticosterone levels only, there were a negative covariation and correlation between intercept and slope (fanning-in pattern), indicating that higher elevations were associated with lower plasticity. For stress-induced levels, the negative covariation was present too, but the strength of correlation was less clear (model 3, table S4).

Lastly, there was a positive correlation between baseline and stress-induced corticosterone levels within-individual and, to some extent, at the population (phenotypic) level (model 4, table S5), indicating that higher baseline corticosterone levels were associated with higher stress-induced levels overall but also within-individual for a given capture (for which both samples for baseline and stress-induced corticosterone levels were collected).

#### 3. Figure

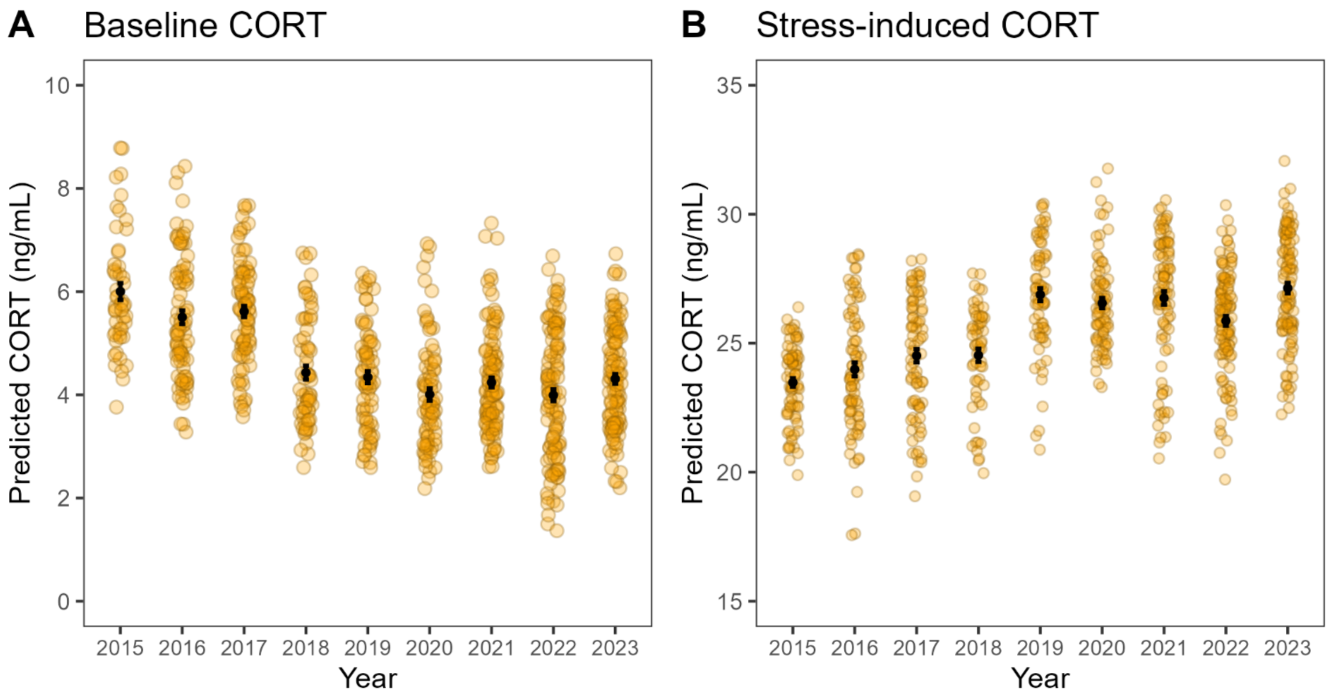

**Figure S1.** Corticosterone levels across years at (A) baseline and (B) stress-induced levels. Yellow circles show individual data predicted from the random regression model (model 3), and black solid circles show means ± SE.

#### 4. Tables

**Table S1. Model 1**: summary statistics of a univariate mixed model fitting baseline or stress-induced corticosterone levels using a Gaussian error distribution as function of sex, year, bleeding time and air temperature at capture. Bird identity was included as random factor. Continuous variables were scaled. Effects are considered relevant when their 95% credible interval (CI) does not overlap zero. Variables of interest for this model are in bold. Baseline corticosterone: N = 783 samples from 417 birds; Stress-induced corticosterone: N = 779 samples from 423 birds.

| | Baseline corticosterone | | | Stress-induced corticosterone | | |
|---|---|---|---|---|---|---|
| | **Posterior mode** | **Posterior median** | **95% CI** | **Posterior mode** | **Posterior median** | **95% CI** |
| **Fixed effects** | | | | | | |
| Intercept | 4.422 | 4.413 | [4.069, 4.703] | 25.23 | 25.276 | [24.01, 26.664] |
| **Sex (female)** | 0.651 | 0.625 | [0.216, 1.048] | 1.334 | 1.189 | [-0.655, 2.938] |
| **Year** | -0.693 | -0.629 | [-0.839, -0.432] | 1.729 | 1.475 | [0.6, 2.373] |
| **Bleeding time** | 1.101 | 1.059 | [0.867, 1.233] | -0.166 | -0.097 | [-0.901, 0.677] |
| **Air temperature** | -0.436 | -0.446 | [-0.627, -0.257] | -2.328 | -2.303 | [-3.132, -1.572] |
| **Random effects** | | | | | | |
| Bird identity | 0.861 | 0.803 | [0, 1.562] | 30.689 | 31.447 | [17.901, 46.589] |
| Residual | 6.792 | 6.904 | [5.915, 7.826] | 94.405 | 98.831 | [84.978, 113.394] |

**Table S2. Model 2**: summary statistics of a univariate mixed model fitting baseline or stress-induced corticosterone levels using a Gaussian error distribution to test between- and within-individual effects of temperature, following a within-subject centring approach. Effects are considered relevant when their 95% credible intervals (CI) do not overlap zero. Variables of interest for this model are in bold. N = 783 samples from 417 birds; Stress-induced corticosterone: N = 779 samples from 423 birds.

| | **Baseline corticosterone** | | | **Stress-induced corticosterone** | | |
|---|---|---|---|---|---|---|
| | **Posterior mode** | **Posterior median** | **95% CI** | **Posterior mode** | **Posterior median** | **95% CI** |
| **Fixed effects** | | | | | | |
| Intercept | 6.112 | 5.904 | [4.909, 7.01] | 34.589 | 34.256 | [29.513, 38.606] |
| Sex (female) | 0.697 | 0.623 | [0.155, 1.007] | 1.085 | 1.181 | [-0.912, 2.865] |
| Year | -0.575 | -0.633 | [-0.846, -0.462] | 1.534 | 1.486 | [0.516, 2.309] |
| Bleeding time | 1.016 | 1.057 | [0.884, 1.255] | -0.095 | -0.078 | [-0.92, 0.652] |
| **Within-ID temperature effect** | -0.11 | -0.108 | [-0.175, -0.054] | -0.423 | -0.521 | [-0.782, -0.276] |
| **Between-ID temperature effect** | -0.094 | -0.106 | [-0.173, -0.031] | -0.679 | -0.636 | [-0.952, -0.342] |
| **Difference of between-within ID effects** | 0.016 | 0.004 | [-0.088, 0.099] | -0.093 | -0.114 | [-0.479, 0.302] |
| **Random effects** | | | | | | |
| Bird identity | 0.922 | 0.84 | [0, 1.547] | 31.211 | 31.214 | [16.546, 44.128] |
| Residual | 6.981 | 6.901 | [6.022, 7.933] | 96.029 | 99.265 | [86.397, 114.305] |

**Table S3. Model comparisons** of a series of univariate mixed models including the same fixed effect structure, including a random intercept or random intercept and slope for temperature and a homogeneous or heterogeneous residual structure.

| **Trait** | **Random effects** | **Residuals** | **DIC** | **ΔDIC** |
|---|---|---|---|---|
| Baseline corticosterone | intercept and slope | heterogeneous | 3657.132 | 0 |
| | intercept | heterogeneous | 3686.434 | 29.302 |
| | intercept and slope | homogeneous | 3778.968 | 121.836 |
| | intercept | homogeneous | 3811.308 | 154.176 |
| Stress-induced corticosterone | intercept and slope | heterogeneous | 5872.668 | 0 |
| | intercept | heterogeneous | 5877.706 | 5.038 |
| | intercept and slope | homogeneous | 5941.948 | 69.280 |
| | intercept | homogeneous | 5943.453 | 70.785 |

**Table S4. Model 3**: summary statistics of a univariate random regression model of baseline or stress-induced corticosterone levels using a Gaussian error distribution, including bird identity as random intercept, air temperature at capture as random slope and a heterogeneous residual structure (one residual per year). Effects are considered relevant when their 95% credible intervals (CI) do not overlap zero. Variables of interest for this model are in bold. Cov. = covariance, corr. = correlation. N = 783 samples from 417 birds; Stress-induced corticosterone: N = 779 samples from 423 birds.

| | **Baseline corticosterone** | | | **Stress-induced corticosterone** | | |
|---|---|---|---|---|---|---|
| | **Posterior mode** | **Posterior median** | **95% CI** | **Posterior mode** | **Posterior median** | **95% CI** |
| **Fixed effects** | | | | | | |
| Intercept | 4.355 | 4.35 | [4.065, 4.624] | 24.96 | 25.119 | [23.875, 26.568] |
| Sex (female) | 0.605 | 0.522 | [0.149, 0.9] | 1.011 | 0.967 | [-0.821, 2.711] |
| Year | -0.572 | -0.5 | [-0.699, -0.284] | 1.293 | 1.458 | [0.452, 2.322] |
| Bleeding time | 1.043 | 1.065 | [0.884, 1.235] | -0.28 | -0.136 | [-0.894, 0.69] |
| Air temperature | -0.405 | -0.371 | [-0.548, -0.186] | -2.028 | -2.173 | [-2.856, -1.369] |
| **Random effects** | | | | | | |
| **Individual intercept** | 0.999 | 1.035 | [0.492, 1.698] | 32.035 | 36.964 | [23.81, 48.978] |
| **Individual slope** | 0.191 | 0.281 | [0, 0.62] | 0.031 | 0.479 | [0, 4.09] |
| **Intercept-slope cov.** | -0.31 | -0.321 | [-0.598, -0.012] | -0.002 | -0.998 | [-6.126, 2.018] |
| **Individual-slope corr.** | -0.603 | -0.641 | [-0.971, -0.249] | -0.484 | -0.292 | [-0.951, 0.554] |
| Residual 1 | 25.638 | 27.409 | [17.86, 39.47] | 118.053 | 122.062 | [77.942, 174.659] |
| Residual 2 | 6.725 | 6.723 | [4.207, 9.454] | 125.032 | 130.071 | [85.116, 181.195] |
| Residual 3 | 6.579 | 6.274 | [4.224, 8.72] | 74.041 | 75.107 | [47.798, 110.275] |
| Residual 4 | 7.191 | 7.776 | [5.298, 11.397] | 150.339 | 154.116 | [90.366, 225.2] |
| Residual 5 | 2.136 | 2.149 | [1.044, 3.354] | 31.357 | 39.322 | [20.205, 64.384] |
| Residual 6 | 5.504 | 5.646 | [3.786, 7.631] | 106.621 | 117.047 | [81.763, 172.869] |
| Residual 7 | 5.319 | 5.554 | [3.88, 7.448] | 82.821 | 79.836 | [49.563, 109.156] |
| Residual 8 | 3.497 | 3.51 | [2.42, 4.736] | 42.184 | 39.289 | [23.686, 59.767] |
| Residual 9 | 2.725 | 2.811 | [1.784, 4.115] | 113.515 | 116.333 | [80.328, 156.091] |

**Table S5. Model 4**: summary statistics of bivariate mixed model to test the covariance and correlation of baseline and stress-induced corticosterone levels. Effects are considered relevant when their 95% credible intervals (CI) do not overlap zero. Variables of interest for this model are in bold. N = 733 samples from 404 birds for both baseline (C0) and stress-induced (C30) corticosterone levels.

| **Fixed effects** | **Response** | **Posterior mode** | **Posterior median** | **95% CI** |
|---|---|---|---|---|
| Intercept | C0 | 4.437 | 4.387 | [4.059, 4.669] |
| | C30 | 25.497 | 25.393 | [24.06, 26.707] |
| Sex (female) | C0 | 0.747 | 0.66 | [0.22, 1.069] |
| | C30 | 0.736 | 0.859 | [-0.788, 2.812] |
| Year | C0 | -0.687 | -0.643 | [-0.854, -0.429] |
| | C30 | 1.478 | 1.42 | [0.579, 2.247] |
| Bleeding time | C0 | 1.024 | 1.041 | [0.855, 1.239] |
| | C30 | -0.189 | -0.315 | [-1.095, 0.443] |
| Air temperature | C0 | -0.408 | -0.444 | [-0.66, -0.256] |
| | C30 | -2.612 | -2.477 | [-3.262, -1.65] |
| **Random effects** | | | | |
| Intercept variance | C0 | 0.963 | 0.85 | [0, 1.599] |
| | C30 | 29.497 | 27.54 | [15.593, 43.517] |
| **Intercept-intercept covariance** | C0-C30 | 1.534 | 1.787 | [-0.253, 4.268] |
| **Intercept-intercept correlation** | C0-C30 | 0.396 | 0.391 | [-0.051, 0.809] |
| Within-ID variance | C0 | 7.033 | 6.832 | [5.794, 7.867] |
| | C30 | 96.23 | 98.58 | [85.946, 114.72] |
| **Within-ID covariance** | C0-C30 | 7.094 | 6.843 | [4.291, 9.639] |
| **Within-ID correlation** | C0-C30 | 0.273 | 0.264 | [0.16, 0.339] |

**Table S6. Model 5b**: summary statistics of a bivariate mixed model to test for the covariance and correlation between baseline or stress-induced corticosterone levels and fledgling number **in males**. Hormonal levels were fitted using a Gaussian error distribution and reproductive success using a zero-inflated Poisson distribution. Therefore, reproductive success shows estimates for the Poisson and zero-inflated (ZI) part of the distribution. Effects are considered relevant when their 95% credible intervals (CI) do not overlap zero. Variables of interest for this model are in bold. Baseline corticosterone: N = 366 samples from 195 birds; Stress-induced corticosterone: N = 368 samples from 200 birds.

| | | **Baseline corticosterone** | | | **Stress-induced corticosterone** | | |
|---|---|---|---|---|---|---|---|
| | | **Posterior mode** | **Posterior median** | **95% CI** | **Posterior mode** | **Posterior median** | **95% CI** |
| **Fixed effects** | **Response** | | | | | | |
| Intercept | CORT | 4.346 | 4.386 | [4.132, 4.651] | 25.46 | 25.537 | [24.293, 26.951] |
| | Fledglings | 1.558 | 1.549 | [1.493, 1.607] | 1.559 | 1.554 | [1.494, 1.609] |
| | Fledglings (ZI) | -3.024 | -3.118 | [-4.149, -2.278] | -2.782 | -3.007 | [-4.037, -2.412] |
| Year | CORT | -0.314 | -0.336 | [-0.619, -0.085] | 2.345 | 2.394 | [1.244, 3.718] |
| | Fledglings | 0.034 | 0.028 | [-0.033, 0.084] | 0.036 | 0.034 | [-0.019, 0.089] |
| | Fledglings (ZI) | -0.159 | -0.183 | [-0.786, 0.417] | -0.082 | -0.119 | [-0.696, 0.406] |
| Bleeding time | CORT | 1.081 | 1.077 | [0.827, 1.302] | 0.06 | -0.063 | [-1.228, 0.969] |
| Air temperature | CORT | -0.576 | -0.561 | [-0.85, -0.299] | -2.273 | -2.596 | [-3.904, -1.473] |
| **Random effects** | | | | | | | |
| Intercept variance | CORT | 0.817 | 0.931 | [0.11, 1.799] | 24.289 | 24.438 | [9.128, 47.01] |
| | Fledglings | 0 | 0.001 | [0, 0.01] | 0 | 0.001 | [0, 0.008] |
| | Fledglings (ZI) | 2.529 | 3.636 | [0, 11.842] | 0.043 | 2.621 | [0, 9.413] |
| Slope variance | CORT slope | 0.011 | 0.416 | [0, 1.238] | 0.039 | 1.404 | [0, 8.834] |
| **Intercept-intercept cov.** | **CORT-fledglings** | 0 | 0.003 | [-0.047, 0.069] | -0.001 | 0.01 | [-0.202, 0.29] |
| **Intercept-intercept corr.** | | 0.891 | 0.226 | [-0.864, 0.997] | 0.35 | 0.142 | [-0.829, 0.963] |
| **Intercept-intercept cov.** | **CORT-fledglings (ZI)** | 0.922 | 0.687 | [-0.279, 1.88] | -0.073 | -1.57 | [-6.828, 3.431] |
| **Intercept-intercept corr.** | | 0.398 | 0.425 | [-0.131, 0.959] | -0.062 | -0.244 | [-0.935, 0.454] |
| Intercept-intercept cov. | Fledglings-fledglings (ZI) | 0 | 0.001 | [-0.124, 0.101] | 0 | -0.003 | [-0.128, 0.078] |
| Intercept-intercept corr. | | -0.584 | 0.019 | [-0.997, 0.82] | -0.854 | -0.212 | [-0.973, 0.793] |
| Intercept-slope cov. | CORT slope-CORT | -0.499 | -0.374 | [-0.909, 0.035] | 0.004 | -1.185 | [-7.569, 3.844] |
| Intercept-slope corr. | | -0.929 | -0.697 | [-0.999, -0.018] | -0.185 | -0.261 | [-0.984, 0.597] |
| **Intercept-slope cov.** | **CORT slope-fledglings** | 0 | -0.001 | [-0.045, 0.034] | 0.001 | -0.001 | [-0.094, 0.076] |
| **Intercept-slope corr.** | | -0.969 | -0.159 | [-0.985, 0.874] | -0.7 | -0.113 | [-0.941, 0.893] |
| **Intercept-slope cov.** | **CORT slope-fledglings (ZI)** | -0.307 | -0.206 | [-1.213, 0.75] | 0.002 | 0.586 | [-1.29, 4.431] |
| **Intercept-slope corr.** | | -0.027 | -0.238 | [-0.985, 0.44] | 0.727 | 0.435 | [-0.65, 0.996] |
| Residual (within-ID) variance | CORT | 5.051 | 5.194 | [4.129, 6.424] | 106.527 | 106.294 | [84.859, 127.151] |
| | Fledglings | 0.039 | 0.041 | [0.026, 0.061] | 0.039 | 0.04 | [0.025, 0.058] |
| | Fledglings (ZI) | 5 | 5 | [5, 5] | 5 | 5 | [5, 5] |

**Table S7. Model 5c**: summary statistics of a bivariate mixed model to test for the covariance and correlation between baseline or stress-induced corticosterone levels and fledgling number **in females**. Hormonal levels were fitted using a Gaussian error distribution and reproductive success using a zero-inflated Poisson distribution. Therefore, reproductive success shows estimates for the Poisson and zero-inflated (ZI) part of the distribution. Effects are considered relevant when their 95% credible intervals (CI) do not overlap zero. Variables of interest for this model are in bold. Baseline corticosterone: N = 394 samples from 211 birds; Stress-induced corticosterone: N = 375 samples from 208 birds.

| | | **Baseline corticosterone** | | | **Stress-induced corticosterone** | | |
|---|---|---|---|---|---|---|---|
| | | **Posterior mode** | **Posterior median** | **95% CI** | **Posterior mode** | **Posterior median** | **95% CI** |
| **Fixed effects** | **Response** | | | | | | |
| Intercept | CORT | 4.983 | 4.987 | [4.697, 5.277] | 26.295 | 26.491 | [25.336, 27.987] |
| | Fledglings | 1.522 | 1.517 | [1.454, 1.577] | 1.529 | 1.526 | [1.464, 1.58] |
| | Fledglings (ZI) | -2.587 | -2.616 | [-3.431, -2.08] | -2.989 | -3.018 | [-3.831, -2.296] |
| Year | CORT | -0.724 | -0.736 | [-1.035, -0.398] | 0.11 | 0.203 | [-1.019, 1.586] |
| | Fledglings | 0.04 | 0.044 | [-0.016, 0.102] | 0.038 | 0.04 | [-0.024, 0.091] |
| | Fledglings (ZI) | -0.236 | -0.234 | [-0.719, 0.301] | -0.332 | -0.317 | [-0.892, 0.22] |
| Bleeding time | CORT | 0.986 | 0.977 | [0.676, 1.247] | 0.161 | -0.076 | [-1.317, 1.036] |
| Air temperature | CORT | -0.417 | -0.397 | [-0.716, -0.141] | -2.498 | -2.418 | [-3.523, -1.354] |
| **Random effects** | | | | | | | |
| Intercept variance | CORT | 1.169 | 0.981 | [0, 1.997] | 38.873 | 43.794 | [20.039, 69.831] |
| | Fledglings | 0 | 0.001 | [0, 0.011] | 0 | 0.001 | [0, 0.009] |
| | Fledglings (ZI) | 0.048 | 2.087 | [0, 7.175] | 3.603 | 3.479 | [0.003, 9.279] |
| Slope variance | CORT slope | 0.006 | 0.325 | [0, 1.105] | 0.028 | 0.947 | [0, 6.808] |
| **Intercept-intercept cov.** | **CORT-fledglings** | 0 | 0.008 | [-0.037, 0.08] | 0.003 | 0.034 | [-0.222, 0.406] |
| **Intercept-intercept corr.** | | 0.904 | 0.409 | [-0.822, 0.999] | 0.493 | 0.257 | [-0.789, 0.977] |
| **Intercept-intercept cov.** | **CORT-fledglings (ZI)** | -0.017 | -0.002 | [-1.093, 1.151] | -3.8 | -4.57 | [-11.259, 1.261] |
| **Intercept-intercept corr.** | | 0.105 | -0.004 | [-0.83, 0.9] | -0.373 | -0.4 | [-0.938, 0.023] |
| Intercept-intercept cov. | Fledglings-fledglings (ZI) | 0 | 0 | [-0.107, 0.089] | 0 | -0.004 | [-0.118, 0.108] |
| Intercept-intercept corr. | | 0.071 | -0.017 | [-0.953, 0.861] | -0.368 | -0.185 | [-0.973, 0.819] |
| Intercept-slope cov. | CORT slope-CORT | -0.001 | -0.403 | [-1.061, 0.074] | 0.01 | -1.454 | [-10.05, 4.051] |
| Intercept-slope corr. | | -0.919 | -0.755 | [-0.997, 0.261] | -0.523 | -0.326 | [-0.997, 0.697] |
| **Intercept-slope cov.** | **CORT slope-fledglings** | 0 | -0.004 | [-0.059, 0.026] | 0 | -0.001 | [-0.094, 0.059] |
| **Intercept-slope corr.** | | -0.879 | -0.409 | [-0.993, 0.871] | 0.044 | -0.107 | [-0.965, 0.882] |
| **Intercept-slope cov.** | **CORT slope-fledglings (ZI)** | 0.008 | 0.004 | [-0.838, 0.778] | 0.002 | 0.425 | [-1.369, 3.559] |
| **Intercept-slope corr.** | | 0.024 | 0.013 | [-0.967, 0.807] | 0.763 | 0.358 | [-0.757, 0.989] |
| Residual (within-ID) variance | CORT | 6.872 | 6.932 | [5.425, 8.553] | 84.678 | 84.805 | [66.775, 104.351] |
| | Fledglings | 0.039 | 0.042 | [0.028, 0.064] | 0.042 | 0.043 | [0.027, 0.063] |
| | Fledglings (ZI) | 5 | 5 | [5, 5] | 5 | 5 | [5, 5] |

**Table S8. Model 6**: summary statistics of bivariate mixed model to test the covariance and correlation of baseline or stress-induced corticosterone levels and **nest outcome (success or failure)**. Hormonal levels were fitted using a Gaussian error distribution and nest outcome using a binomial distribution. Effects are considered relevant when their 95% credible intervals (CI) do not overlap zero. Variables of interest for this model are in bold. Cov. = covariance, corr. = correlation, Int. = intercept. Baseline corticosterone: N = 760 samples from 406 birds; Stress-induced corticosterone: N = 743 samples from 408 birds.

| | | **Baseline corticosterone** | | | **Stress-induced corticosterone** | | |
|---|---|---|---|---|---|---|---|
| **Fixed effects** | **Response** | **Posterior mode** | **Posterior median** | **95% CI** | **Posterior mode** | **Posterior median** | **95% CI** |
| Intercept | CORT | 4.373 | 4.383 | [4.078, 4.702] | 25.486 | 25.487 | [24.317, 26.89] |
| | Nest outcome (failure) | -2.026 | -2.004 | [-2.49, -1.6] | -2.093 | -2.03 | [-2.539, -1.606] |
| Sex (female) | CORT | 0.596 | 0.617 | [0.224, 1.054] | 1.219 | 1.041 | [-0.869, 2.762] |
| | Nest outcome (failure) | 0.272 | 0.229 | [-0.254, 0.757] | 0.063 | 0.084 | [-0.487, 0.567] |
| Year | CORT | -0.511 | -0.528 | [-0.722, -0.322] | 1.487 | 1.416 | [0.45, 2.291] |
| | Nest outcome (failure) | -0.187 | -0.146 | [-0.392, 0.124] | -0.157 | -0.17 | [-0.46, 0.06] |
| Bleeding time | CORT | 1.012 | 1.024 | [0.844, 1.212] | 0.064 | -0.022 | [-0.831, 0.786] |
| Air temperature | CORT | -0.463 | -0.484 | [-0.679, -0.284] | -2.521 | -2.553 | [-3.369, -1.669] |
| **Random effects** | | | | | | | |
| Intercept variance | CORT | 1.123 | 1.098 | [0.473, 1.777] | 27.394 | 30.956 | [17, 46.459] |
| | Nest outcome | 0.923 | 1.319 | [0, 2.894] | 1.094 | 1.365 | [0.113, 2.975] |
| Slope variance | CORT slope | 0.317 | 0.409 | [0.042, 0.917] | 0.028 | 0.849 | [0, 4.767] |
| **Int.-int. cov.** | **CORT-nest outcome** | 0.211 | 0.214 | [-0.296, 0.671] | -2.211 | -2.028 | [-4.662, 0.181] |
| **Int.-int. corr.** | | 0.222 | 0.196 | [-0.238, 0.711] | -0.352 | -0.333 | [-0.74, 0.034] |
| Int.-slope cov. | CORT slope-CORT | -0.535 | -0.514 | [-0.933, -0.148] | -0.169 | -1.257 | [-6.119, 2.509] |
| Int.-slope corr. | | -0.782 | -0.788 | [-0.993, -0.469] | -0.318 | -0.307 | [-0.874, 0.505] |
| **Int.-slope cov.** | **CORT slope-nest outcome** | -0.104 | -0.099 | [-0.455, 0.258] | 0.018 | 0.338 | [-0.364, 1.973] |
| **Int.-slope corr.** | | -0.252 | -0.156 | [-0.799, 0.341] | 0.635 | 0.396 | [-0.409, 0.975] |
| Residual (within-ID) variance | CORT | 5.917 | 6.007 | [5.121, 6.848] | 100.165 | 99.373 | [85.959, 115.22] |
| | Nest outcome | 1 | 1 | [1, 1] | 1 | 1 | [1, 1] |

**Table S9. Model 7**: summary statistics of bivariate mixed model to test the covariance and correlation of baseline or stress-induced corticosterone levels and **reproductive success (fledgling number) in successful nests only**. Hormonal levels were fitted using a Gaussian error distribution and reproductive success using a Poisson error distribution. Effects are considered relevant when their 95% credible intervals (CI) do not overlap zero. Variables of interest for this model are in bold. Cov. = covariance, corr. = correlation, Int. = intercept. Baseline corticosterone: N = 607 samples from 348 birds; Stress-induced corticosterone: N = 601 samples from 354 birds.

| | | **Baseline corticosterone** | | | **Stress-induced corticosterone** | | |
|---|---|---|---|---|---|---|---|
| **Fixed effect** | **Response** | **Posterior mode** | **Posterior median** | **95% CI** | **Posterior mode** | **Posterior median** | **95% CI** |
| Intercept | CORT | 4.449 | 4.348 | [4.008, 4.668] | 25.498 | 25.717 | [24.304, 27.161] |
| | Fledglings | 1.557 | 1.56 | [1.499, 1.62] | 1.576 | 1.565 | [1.506, 1.621] |
| Sex (female) | CORT | 0.629 | 0.563 | [0.15, 1.024] | 0.963 | 1.106 | [-0.911, 3.02] |
| | Fledglings | -0.027 | -0.031 | [-0.112, 0.052] | -0.028 | -0.027 | [-0.108, 0.06] |
| Year | CORT | -0.601 | -0.553 | [-0.775, -0.336] | 0.962 | 1.044 | [0.152, 2.044] |
| | Fledglings | 0.033 | 0.036 | [-0.007, 0.07] | 0.027 | 0.036 | [-0.006, 0.074] |
| Bleeding time | CORT | 0.926 | 0.915 | [0.721, 1.137] | 0.242 | 0.066 | [-0.829, 0.883] |
| Air temperature | CORT | -0.446 | -0.407 | [-0.607, -0.169] | -2.748 | -2.712 | [-3.592, -1.761] |
| **Random effects** | | | | | | | |
| Intercept variance | CORT | 1.042 | 1.216 | [0.512, 2.185] | 31.41 | 34.715 | [18.615, 52.497] |
| | Fledglings | 0 | 0.001 | [0, 0.005] | 0 | 0 | [0, 0.004] |
| Slope variance | CORT slope | 0.364 | 0.344 | [0, 0.884] | 0.034 | 0.946 | [0, 5.26] |
| **Int.-int. cov.** | **CORT-Fledglings** | 0 | 0.01 | [-0.021, 0.068] | 0 | 0.011 | [-0.125, 0.261] |
| **Int.-int. corr.** | | 0.775 | 0.481 | [-0.712, 0.988] | 0.52 | 0.156 | [-0.701, 0.956] |
| Int.-slope cov. | CORT slope-CORT | -0.422 | -0.475 | [-0.935, 0] | -0.044 | -2.066 | [-9.29, 1.535] |
| Int.-slope corr. | | -0.909 | -0.78 | [-0.989, -0.318] | -0.737 | -0.428 | [-0.954, 0.394] |
| **Int.-slope cov.** | **CORT slope-fledglings** | 0 | -0.005 | [-0.041, 0.014] | 0 | 0 | [-0.047, 0.037] |
| **Int.-slope corr.** | | -0.907 | -0.486 | [-0.98, 0.737] | 0.1 | -0.098 | [-0.914, 0.807] |
| Residual (within-ID) variance | CORT | 6.117 | 5.829 | [4.895, 6.945] | 91.009 | 94.256 | [77.833, 110.254] |
| | Fledglings | 0.051 | 0.05 | [0.039, 0.063] | 0.048 | 0.05 | [0.038, 0.064] |

**Table S10. Model 8:** summary statistics of bivariate mixed model to test the covariance and correlation of baseline or stress-induced corticosterone levels and **reproductive success (brood mass) in successful nests only**. Hormonal levels were fitted using a Gaussian error distribution and reproductive success using a Poisson error distribution. Effects are considered relevant when their 95% credible intervals (CI) do not overlap zero. Variables of interest for this model are in bold. Cov. = covariance, corr. = correlation, Int. = intercept. Baseline corticosterone: N = 538 samples from 326 birds; Stress-induced corticosterone: N = 534 samples from 332 birds.

| | | **Baseline corticosterone** | | | **Stress-induced corticosterone** | | |
|---|---|---|---|---|---|---|---|
| **Fixed effect** | **Response** | **Posterior mode** | **Posterior median** | **95% CI** | **Posterior mode** | **Posterior median** | **95% CI** |
| Intercept | CORT | 4.239 | 4.3 | [3.908, 4.619] | 25.911 | 26.069 | [24.396, 27.482] |
| | Brood mass | 79.842 | 80.021 | [76.499, 83.466] | 80.114 | 80.194 | [77.324, 83.995] |
| Sex (female) | CORT | 0.55 | 0.619 | [0.093, 1.109] | 1.117 | 1.196 | [-0.966, 3.519] |
| | Brood mass | -2.883 | -2.112 | [-6.805, 2.775] | -1.155 | -1.357 | [-6.41, 3.335] |
| Year | CORT | -0.586 | -0.629 | [-0.849, -0.379] | 0.903 | 0.676 | [-0.478, 1.569] |
| | Brood mass | 2.991 | 2.699 | [0.365, 4.937] | 2.984 | 2.613 | [0.266, 4.739] |
| Bleeding time | CORT | 0.948 | 0.951 | [0.714, 1.153] | 0.083 | 0.075 | [-0.895, 0.999] |
| Air temperature | CORT | -0.369 | -0.372 | [-0.608, -0.122] | -2.538 | -2.651 | [-3.679, -1.614] |
| **Random effects** | | | | | | | |
| Intercept variance | CORT | 1.062 | 1.159 | [0.334, 2.179] | 45.652 | 50.215 | [30.088, 72.167] |
| Intercept variance | Brood mass | 0.859 | 40.47 | [0.002, 114.722] | 0.49 | 47.502 | [0.003, 141.115] |
| Slope variance | CORT slope | 0.003 | 0.314 | [0, 0.969] | 0.074 | 3.552 | [0.001, 12.212] |
| **Int.-int. cov.** | **CORT-Brood mass** | 4.061 | 3.491 | [-0.555, 9.289] | 0.205 | 7.436 | [-14.306, 31.065] |
| **Int.-int. corr.** | | 0.731 | 0.563 | [-0.121, 0.991] | 0.163 | 0.17 | [-0.418, 0.758] |
| Int.-slope cov. | CORT slope-CORT | -0.339 | -0.412 | [-0.912, 0.047] | -7.731 | -6.428 | [-15.274, 0.909] |
| Int.-slope corr. | | -0.836 | -0.728 | [-0.99, -0.112] | -0.58 | -0.517 | [-0.931, -0.049] |
| **Int.-slope cov.** | **CORT slope-Brood mass** | -0.013 | -1.45 | [-5.508, 0.988] | -0.027 | -1.069 | [-11.796, 8.895] |
| **Int.-slope corr.** | | -0.665 | -0.53 | [-0.978, 0.375] | -0.123 | -0.159 | [-0.876, 0.643] |
| Residual (within-ID) variance | CORT | 5.94 | 5.961 | [4.975, 7.161] | 74.712 | 78.687 | [62.732, 95.33] |
| | Brood mass | 725.051 | 727.847 | [621.209, 831.545] | 725.226 | 703.651 | [585.101, 829.609] |
